# A joint alpha power–phase dynamic shapes visual sensitivity

**DOI:** 10.64898/2026.02.19.706926

**Authors:** Henry Beale, Jason Mattingley, Anthony Harris

## Abstract

Perceptual experience is influenced by alpha-band oscillations (8–14 Hz) that dominate parietal and sensory cortices. However, there is uncertainty around the perceptual mechanisms that are affected by oscillatory power and phase. Previous work has linked power and phase to behaviour separately despite their theorised common effects via pulsed-inhibition, potentially misrepresenting or conflating their effects. Here we recorded brain activity using electroencephalography to investigate how alpha oscillations affect the psychometric function over visual contrast in both detection and discrimination tasks. We found that prestimulus power and phase predicted the strength of subsequent evoked neural responses and behavioural accuracy. We then combined power and phase into a joint model of pulsed-inhibition and estimated its effects within a signal detection model of behaviour. The model revealed response gain modulation of visual sensitivity in both tasks, and perceptual bias modulation in detection. Critically, oscillatory power suppressed visual sensitivity more strongly than phase, suggesting a sustained effect of alpha oscillations that is not accounted for by the pulsed-inhibition hypothesis. We conclude that alpha-band activity shapes visual perception by divisively suppressing sensory evidence and baseline sensory noise, with joint power–phase modelling revealing asymmetric contributions to visual sensitivity.

## 1 Introduction

Human visual behaviour relies on inferences about the external world that are shaped by both sensory input and the internal states of the brain’s sensory system. Even when sensory input is held constant, perceptual judgements can vary substantially owing to rhythmic oscillations in the excitability of sensory neural populations [1]. While it is well-accepted that neural oscillations are ubiquitous in sensory cortices and are linked to a variety of cognitive functions (e.g. visual attention, prediction, memory; [2, 3]), their specific impact on the computations underlying visual behaviour remains an open question.

Neural oscillations are a widespread feature of brain activity that are preserved across species [4] and arise from neuronal micro-circuitry that features inhibitory dynamics [5]. Low-frequency oscillations, particularly in the alpha band (8–14 Hz), reflect cycles of cortical excitability that shape the temporal dynamics of neural activity and putatively coordinate information processing amongst brain regions [6–9]. In sensory cortices, neuronal spiking is modulated by both the phase and amplitude of alpha oscillations [10, 11]. Theoretical accounts suggest that alpha oscillations reflect a rhythmic pulsed-inhibition of neural activity (see Fig. 1a) that can be flexibly shaped to influence sensory processing in a goal-dependent manner [12–16].

**Figure 1:**
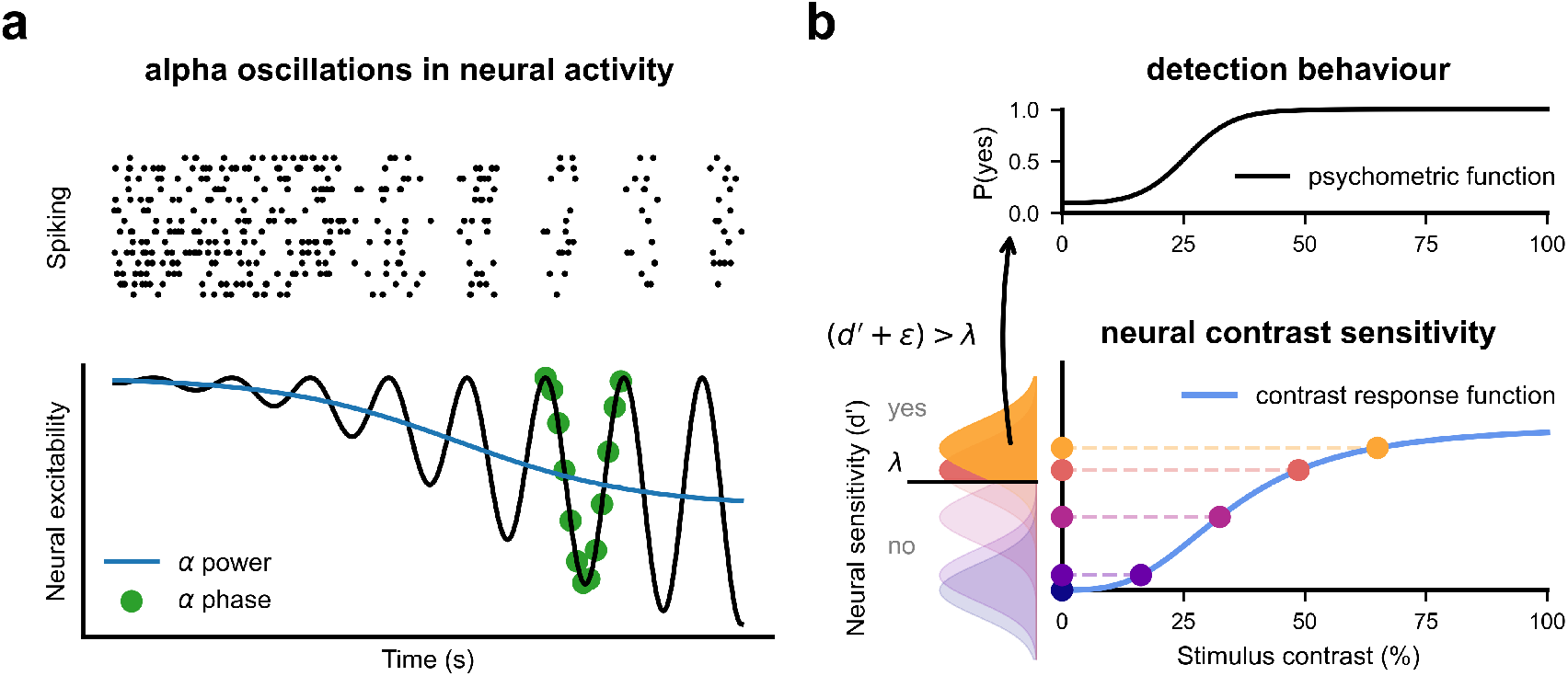
Theoretical relationship between neural activity, alpha oscillations, and perception. **a** Rhythmic suppression of neural spiking activity at ∼10 Hz (upper panel: raster plot showing the firing of a population of simulated neurons). The plot shows rhythmic inhibition emerging gradually toward the right. Stronger pulses of inhibition spaced 100 ms apart result in synchronous firing. Lower panel shows inhibitory pulses (black line) in population level activity reflected in local field potentials or scalp EEG. Pulsed-inhibition underlies measures of alpha-band activity such as power, which increases as a consequence of stronger inhibitory pulses (blue line), and phase (green dots). Adapted from [16]. **b** According to Signal Detection Theory, neural sensitivity to stimulus contrast intensity produces a psychometric function in detection (upper panel; x-axis: contrast; y-axis: probability of detection). While the psychometric function is observed experimentally, performance is determined by latent neural responses (lower panel). The coloured dots show evenly-spaced contrast values that produce neuronal responses of varying strength (due to a nonlinear transducer function in the early visual system; blue line). Neural noise shifts response strength (*d*^*′*^) along the y-axis (lower panel) and produces a distribution across trials. A decision rule asserts that a detection response is made when the neural response (plus noise, *ϵ*) exceeds a criterion level (horizontal black line, *λ*). The psychometric function thus reflects the probability that noisy neural responses exceed this criterion as a function of contrast.

In humans, alpha oscillations are readily measured using non-invasive methods with high temporal resolution, such as electroencephalograpy (EEG) and magnetoencephalography (MEG). The phase of spontaneous alpha oscillations predicts perceptual reports of weak, near-threshold visual stimulation [17–19]. Phasic modulation of visual detection also depends on strong alpha amplitude [20, 21], a relationship supported by causal evidence from Transcranial Magnetic Stimulation (TMS) over occipital cortex [22]. These findings are consistent with a pulsed functional inhibition hypothesis [6, 15] in which the strength of phasic modulation depends on the amplitude of the spontaneous oscillation. That is, inhibitory brain states differ mostly strongly from peak to trough during high amplitude oscillations, and this phasic modulation is diminished during weak oscillations. However, the computational mechanisms by which pulsed inhibition rhythmically modulates perception remain unclear.

Signal Detection Theory (SDT; [23]) provides a powerful tool with which to investigate the computations underlying fluctuations in visual sensitivity. In the context of visual neuroscience, SDT provides a generative model of visual behaviour based on latent neural responses and a decision rule (Fig. 1b). Early studies that linked alpha phase to visual perception did not use SDT methods [17, 21] and thus could not separate perceptual from decisional influences of alpha phase on behaviour, as would be possible with SDT. Recent studies have used SDT to relate changes in alpha power (alpha phase was not measured) to fluctuations in decision bias, rather than sensory sensitivity [24–26]. Critically, however, these studies only assessed performance at a single contrast level. Neural sensitivity is known to vary non-linearly with stimulus intensity [27], meaning that perception could exhibit different patterns of modulation depending on the stimulus intensity range examined. Testing the full range of stimulus intensity values is therefore required to gain a complete picture of how neural oscillations relate to perception.

Distinct computational mechanisms can shape the psychometric function in characteristically different ways, such as the response gain and contrast gain computations that have been proposed for attentional modulation of neural responses (e.g. [28]). Only two studies have sought to relate alpha power to visual sensitivity using a full range of stimulus intensities and yielded separate support for response gain [29] or baseline/criterion modulation [30]. Critically, both studies modelled psychometric functions using detection rates, instead of underlying SDT parameters, and neither examined the association between these mechanisms and the phase of alpha oscillations. A clear understanding of how alpha oscillations shape visual processing, via pulsed inhibition, requires the use of SDT analyses across the complete stimulus intensity range, combined with an analysis of the joint influence of alpha power and phase.

In the present study, we investigated the combined interactive effects of alpha power and phase on visual judgements. We used EEG to measure prestimulus alpha power and phase, and combined these into a modelled inhibitory dynamic that is consistent with the pulsed-inhibition hypothesis of alpha oscillations [12, 15, 16, 31]. We tested observers in both visual detection and discrimination across a wide contrast-intensity range, and we developed a pulsed-inhibition model that links oscillatory phase and power to latent sensory responses using SDT. As described below, alpha-based inhibition modulated the response gain of visual sensitivity in both tasks, whereas a baseline shift in excitability was evident only in detection. These results support a common perceptual mechanism through which alpha power and phase, together, regulate visual sensitivity. Critically, we found that the effect of phase within the pulsed-inhibition model was weaker relative to the power-related suppression of sensory responses, a previously unseen power–phase dynamic that is not captured by pulsed-inhibition models of rhythmic perception.

## 2 Results

### 2.1 Prestimulus alpha oscillations modulate visual responses

To investigate the effects of alpha oscillations on visual perception, we recorded EEG while participants made visual judgements in both detection and discrimination tasks (see Methods). Participants (*n* = 8 x 4 sessions, per task) viewed brief (8 ms) oriented gratings of variable contrast intensities. In the detection task, participants indicated the presence or absence of the target within a set of spatial markers in the visual periphery (see Fig. 2a). In the discrimination task, participants judged whether the stimulus was spatially offset either to the left or right of the markers. Stimulus visibility was manipulated using contrast intensity, which was adaptively set in each session to both probe performance around threshold level and best estimate the full psychometric function over contrast. Behavioural performance improved as a function of stimulus contrast in both tasks, shown by increasing rates of detection (Fig. 2b) and greater accuracy of location discrimination (Fig. 2c) with higher contrast. We used independent component analysis to isolate (per session) a source of posterior scalp EEG activity in the contralateral hemisphere, relative to the stimulus, that responded to stimulus presentation. In both tasks, the magnitude of the stimulus-evoked responses was driven exogenously by stimulus contrast (Fig. 2d) supporting the source components’ relevance for visual processing.

**Figure 2:**
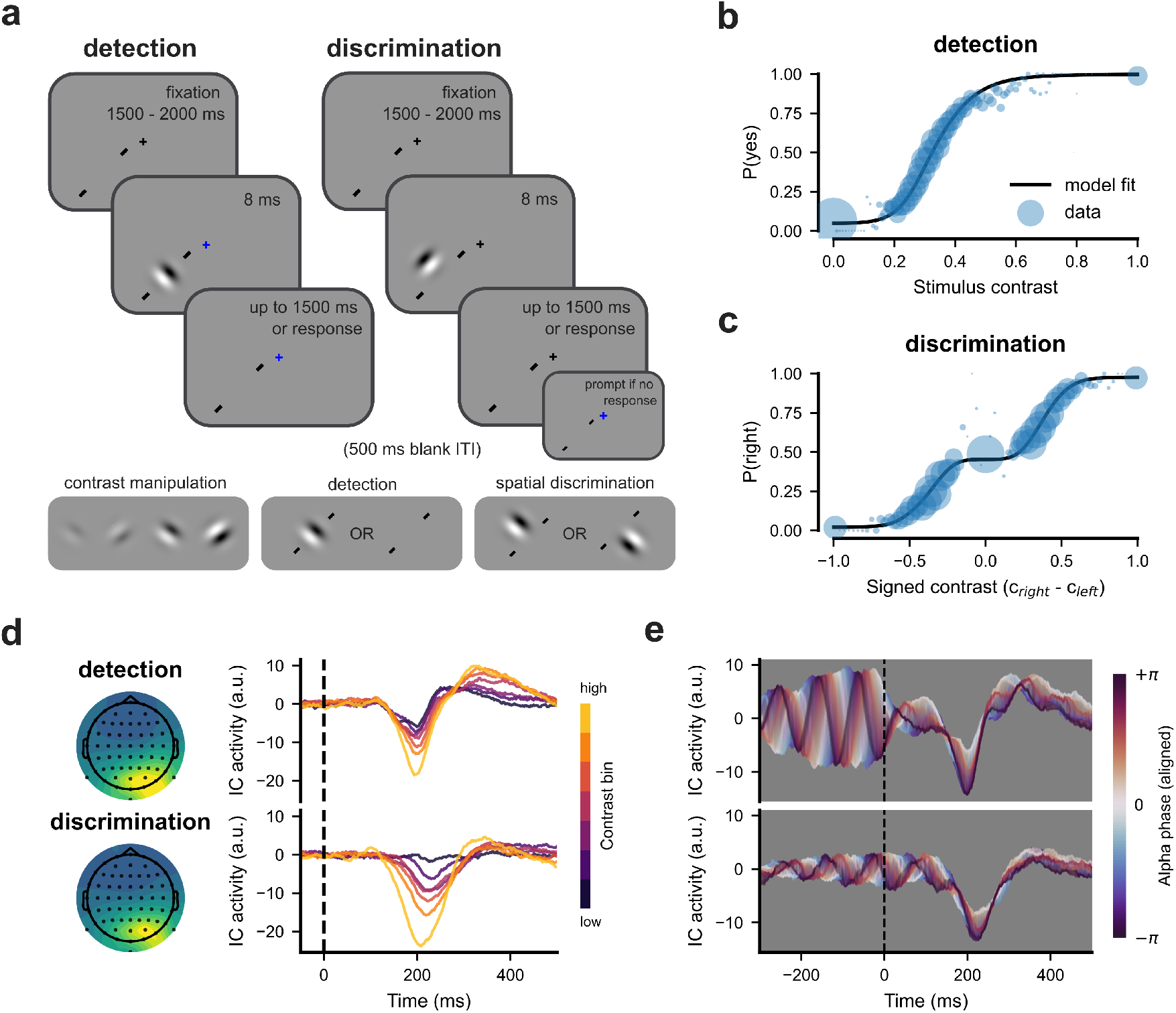
Behaviour and EEG responses in the two visual tasks. **a** Participants viewed grating stimuli that varied in contrast and, in separate experiments, reported stimulus presence/absence (detection; left-hand panels) or spatial position relative to onscreen diagonal tick markers (discrimination; right-hand panels). **b** Psychometric function of contrast-driven detection responses. Black line shows the predicted detection rate from a signal detection model, fitted hierarchically over participants. Circles show data averaged over participants, with sizes reflecting the number of trials presented at each contrast. **c** Psychometric function of spatial discrimination judgements over stimulus contrast (conventions as in **b**). The abscissa shows the signed contrast difference between the right (positive) and left (negative) spatial locations. The ordinate is the probability of reporting the stimulus in the rightward location. **d** Left panel: Scalp topographies of sensor activations from the analysed independent components (z-scored and averaged over sessions; stronger activation is shown brighter, arbitrary units). Right panel: Evoked EEG responses increased with stimulus contrast, brighter lines show increasing levels of binned contrast (dashed line shows stimulus onset time). **e** Evoked responses plotted by alpha-band phase at stimulus onset (estimated a sliding Von Mises window, *σ* = 15*°*, over phase values aligned, per session, to the phase with greatest evoked activation).

We next examined whether the evoked visual responses were modulated endogenously by spontaneous alpha-band activity. Prestimulus oscillations were detected on a single-trial basis using a spectral parameterisation algorithm to separate oscillations from background aperiodic activity [32], using data in a 500 ms window prior to stimulus onset. Estimates of instantaneous power and phase were then obtained for detected oscillations in the 8–14 Hz alpha frequency range. In both tasks, alpha power significantly predicted the amplitude of peaks in the evoked neural activity. Specifically, the negative peaks at ∼200 ms post-stimulus were weaker when alpha power was stronger at stimulus onset (*detection*: *β* = −0.066, *t*_(19345)_ = 2.62, *P* = .009; *discrimination*: *β* = −0.106, *t*_(19345)_ = 2.72, *P* = .007), consistent with previous work showing suppression of early event-related potentials by alpha ongoing oscillations [33]. Furthermore, alpha phase at stimulus onset significantly predicted peak evoked amplitudes in both tasks (*detection*: *d*_*z*_ = 1.287, *P* < .001; *discrimination*: *d*_*z*_ = 1.745, *P* < .001). Fig. 2e shows the evoked waveforms plotted as a function of phase. In summary, the modulation of evoked responses by both power and phase demonstrates that neural responses to the stimuli were modulated endogenously by the state of spontaneous alpha activity at stimulus onset.

### 2.2 Behavioural accuracy modulation by alpha power and phase

Next, we asked whether alpha power and phase impacted perception by modulating behavioural performance across the two tasks. We first used a binning approach to quantify behavioural accuracy and to visualise performance modulation across contrast, shown in fig. 3a–b, consistent with past approaches ([29]; see Methods). Performance was equated across sessions by computing contrast as a percentage of thresholds and behavioural accuracy was then binned by alpha power and phase within each contrast level. To compute a summary measure of performance modulation that aggregates over contrast-specific effects, the slope of alpha-related modulation was averaged across contrast bins. We found stronger alpha power significantly decreased ‘stimulus present’ responses in the detection task (mean slope across contrast bins [95% highest density interval (HDI)] = −0.005 [−.01, −0.001], bootstrapped *P* value (*P*_*boot*_) = .005). Similarly, stronger alpha power led to significantly fewer reports of the correct target location in the discrimination task (mean slope = −0.005 [−0.01, 4e-5], *P*_*boot*_ = .024). Given that prestimulus phase modulated the evoked responses, phase values were circularly realigned relative to the phase that predicted the strongest evoked response. Behavioural responses were then binned by the circular distance from this optimal phase. However, phase did not influence overall responses, collapsed across contrast bins, in either task (*detection*: mean slope = −0.003 [−0.007, 0.002], *P*_*boot*_ = .147; *discrimination*: mean slope = −0.001 [−0.006, 0.005], *P*_*boot*_ = .425). Thus, overall behavioural accuracy was substantially modulated by alpha power, but not by alpha phase.

**Figure 3:**
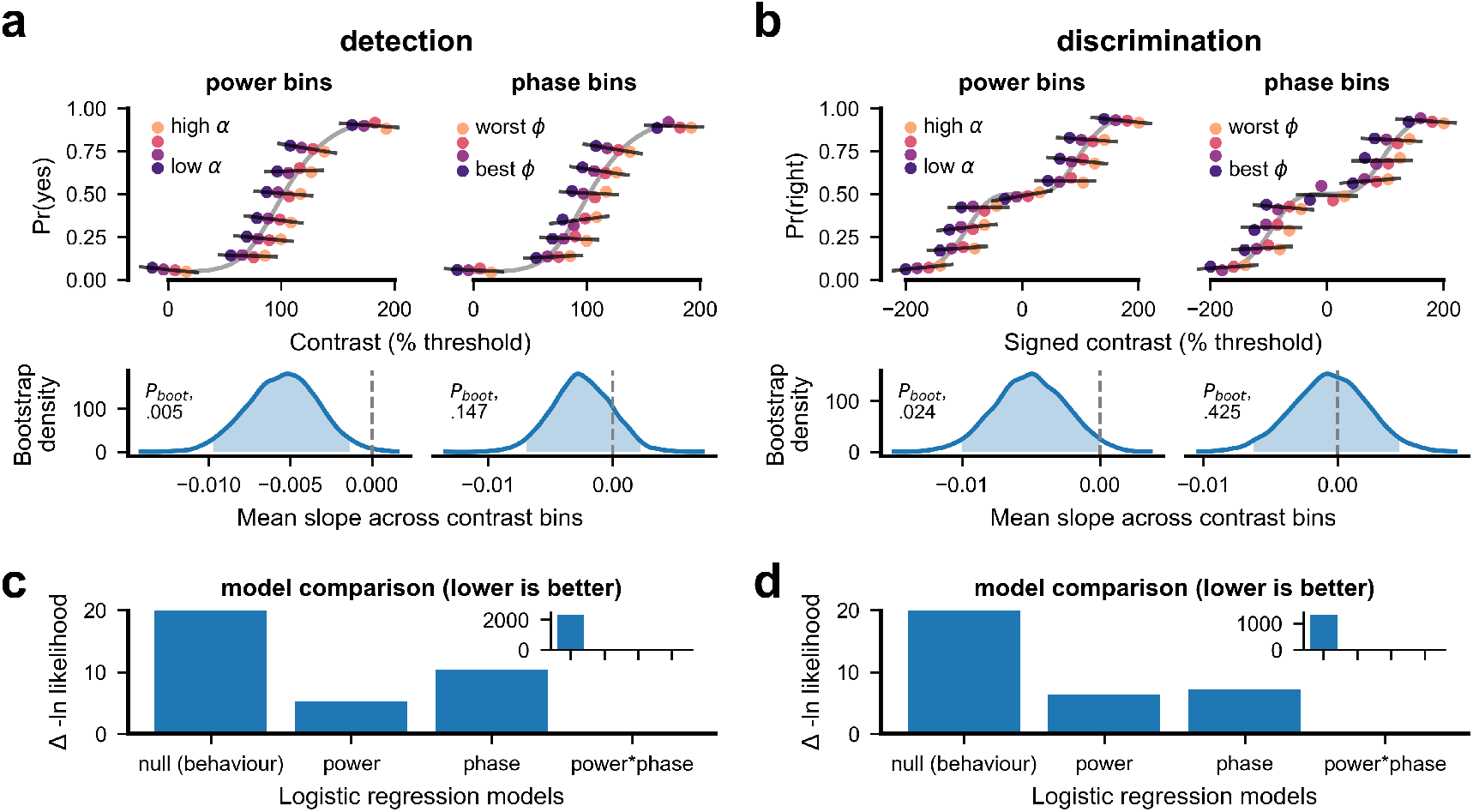
Behavioural accuracy by power and phase. **a–b** Binning analyses of accuracy for the detection (**a**) and discrimination tasks (**b**). Upper panels show accuracy binned by stimulus contrast for power (left) and phase (right). Darker colours show power/phase bins where theory suggests performance should be improved. Contrast jitter (i.e. spreading across the x-axis within each group of points) was added to facilitate visual inspection of power/phase differences within each contrast bin. The dark lines show the fitted slope of accuracy modulation within a contrast bin. Lower panels show the density of the average modulation slope across bins (estimated using kernel density estimation based on 5,000 bootstrap samples). Inset text shows the probability of samples lying below zero (dashed gray line). **c–d** Comparison of logistic regression models fitted to detection (**c**) and discrimination (**d**) data. Separate model fits were conducted for behaviour only (null), power, phase, and power by phase interaction terms. The y-axis (truncated) shows the change in negative log likelihood relative to the best fitting (power^*^phase) model, higher values indicate worse fit. Inset shows the full y-axis which captures the worse fit of the null model.

To investigate how alpha power and phase might modulate behaviour in a way that interacts with stimulus contrast, we conducted logistic regression analyses to predict behavioural accuracy as a function of contrast (see Methods). In the detection task, the addition of both power and phase predictors significantly contributed to the model likelihood relative to a null behaviour-only model (nested likelihood ratio tests; *power* : 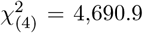, *P* < .001, AIC: 24,175, BIC: 24,248; *phase*: 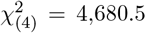, *P* < .001, AIC: 24,185, BIC: 24,258; *null*: AIC: 28,858, BIC: 28,899). This was also true for the discrimination task (*power* : 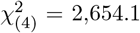, *P* < .001, AIC: 20,081, BIC: 20,151; *phase*: 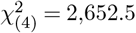, *P* < .001, AIC: 20,082, BIC: 20,153; *null*: AIC: 22,727, BIC: 22,767). The interaction between power and phase added significantly to model likelihood in detection (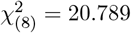, *P* = .008, AIC: 24,180, BIC: 24,318) but not for discrimination (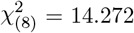, *P* = .075, AIC: 20,084, BIC: 20,218). In both tasks, the best negative log-likelihood was found for the power–phase interaction models (shown in Fig. 3c&d). However, information criteria that include a penalty for model complexity, such as AIC and BIC, consistently favoured the power model more strongly in both tasks (Supplementary Fig. 1). Together, these results suggest that both power and phase modulate visual performance, with an interaction effect emerging in the detection task, but there is stronger evidence for power as a model predictor of behavioural modulation.

### 2.3 Pulsed-inhibition alters response gain and baseline activity

Next, we investigated how alpha oscillations modulate the perceptual mechanisms that give rise to the specific shape of the psychometric function. We used a generative signal detection model to explain behaviour in terms of the latent sensory responses that underlie detection and discrimination choices. The strength of these modelled sensory responses is a function of stimulus contrast (see Fig. 1b) and, we hypothesise, may also be impacted by alpha-related inhibition. Importantly, we combined alpha power and phase estimates into a model of pulsed-inhibition (in line with Fig. 1a; see *Pulsed-inhibition modelling* in the Methods) to compare the computational mechanisms by which inhibition shapes these latent sensory responses, and thus behaviour.

We fitted four models that describe the potential sensory modulation by pulsed-inhibition (see Fig. 4a–b, left columns). First, inhibition might affect visual sensitivity via *contrast gain* which produces poorer sensitivity to weaker contrast values and is reflected in a rightward shift in the psychometric functions. Second, inhibition could divisively scale visual responses, via *response gain*, without changing the underlying range of contrast that the visual system is sensitive to. Third, *noise* in visual responses might be altered by inhibition, which affects performance mostly at the lowest and highest contrast levels. Finally, inhibition might affect the strength of visual responses regardless of contrast, via *criterion* modulation, which affects baseline performance in the psychometric function (i.e. zero-contrast performance in detection, or biased location choice in spatial discrimination).

**Figure 4:**
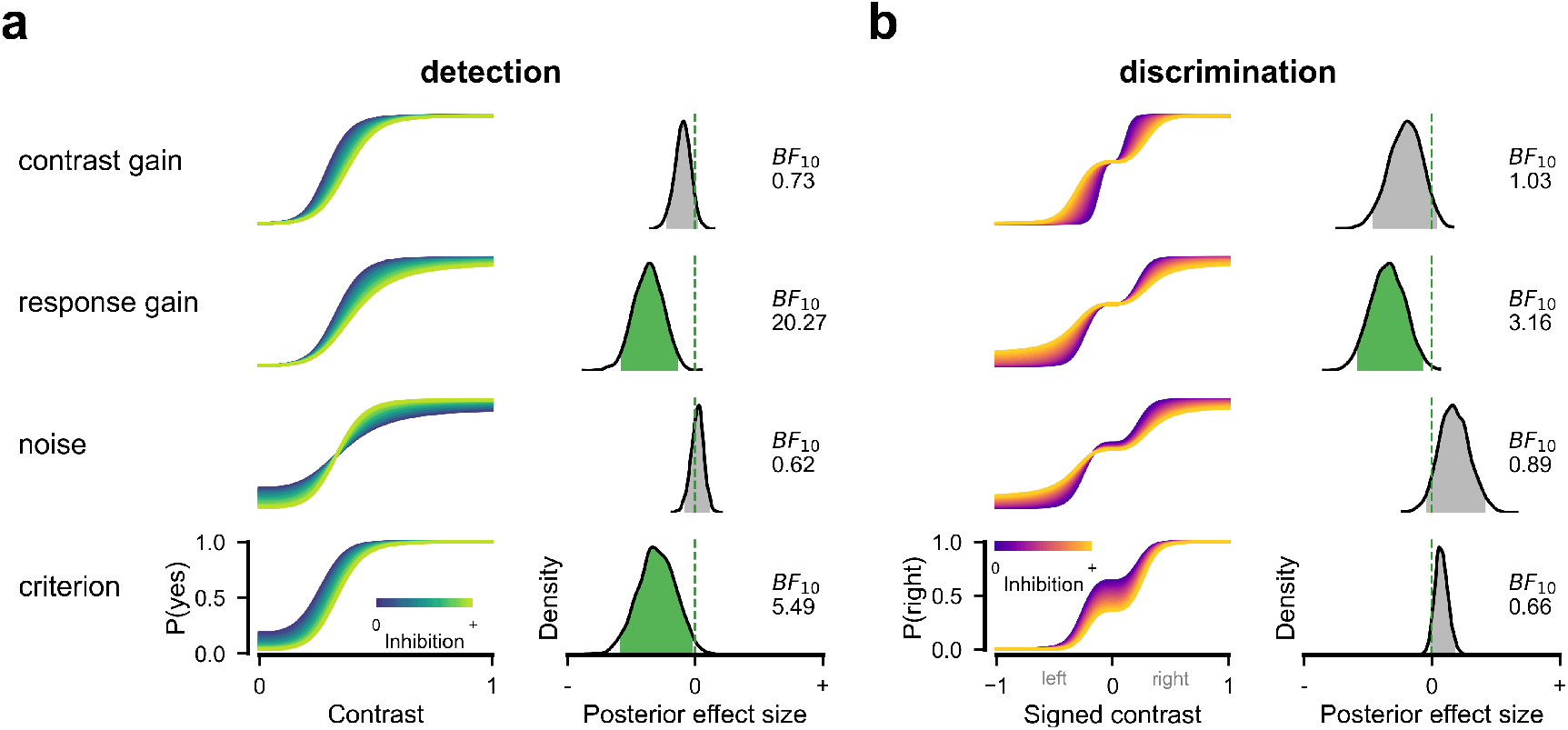
Model-based effects of alpha inhibition on the psychometric function. **a** Signal detection modelling in which pulsed-inhibition modulates the detection psychometric function via four putative mechanisms: contrast gain, response gain, (sensory) noise, and criterion. Left column: Simulated modulations of each parameter in the psychometric function under increasing inhibition (lighter colours). In detection, the y-axis is the probability of a ‘yes’ response. Right column: The inferred posterior densities of the modulation effects. To facilitate comparison, parameter values were standardised using the standard deviation among the hierarchically fitted sessions. Shaded regions of the distribution show the 95% Highest Density Interval and are shown in green when this does not include zero (shown by green vertical dashed line). Text insets show the Bayes Factor in support of the alternative. **b** Inhibition models for the discrimination task. Conventions as in **a** apart from the left column, where simulations of the psychometric function for discrimination are plotted over signed contrast (x-axis) and show the probability of reporting the stimulus as occupying the right-side location (y-axis). The underlying effects within the signal detection models do not change between tasks, but the psychometric function is nonetheless different because of the two-choice response structure.

In the detection task, we found the clearest support for pulsed-inhibition of sensory response gain (Fig. 4a; standardised parameter estimate (*β*_*z*_ [95% HDI]) = −3.388 [−5.413, −1.253], Bayes Factor (BF_10_) = 20.27). Additional support was found for pulsed-inhibition of criterion (*β*_*z*_ = −2.814 [−5.497, −0.191], BF_10_ = 5.49), in line with previous findings [30]. We found no support for oscillations affecting either contrast gain (*β*_*z*_ = −0.899 [−2.113, 0.227], BF_10_ = 0.73) or sensory noise (*β*_*z*_ = 0.167 [−0.79, 1.087], BF_10_ = 0.62). In the discrimination task, we found additional support for response gain modulation (*β*_*z*_ = −2.499 [−4.256, −0.48], BF_10_ = 3.16), but not for any of the remaining models (*contrast gain*: *β*_*z*_ = −1.509 [−3.382, 0.3], BF_10_ = 1.03; *noise*: *β*_*z*_ = 1.283 [−0.325, 3.07], BF_10_ = 0.89; *criterion*: *β*_*z*_ = 0.556 [−0.127, 1.339], BF_10_ = 0.66). Together, the modelling results suggest that, across tasks, alpha inhibits visual sensitivity to contrast via response gain modulation, and via baseline neural activity in the detection task but not in the discrimination task. We note the latter result is explained by the two-alternative forced-choice design of the discrimination task, which forces changes in perceptual bias to be matched between choice options and to thereby cancel out.

### 2.4 Perception is modulated by both sustained and pulsed inhibition

We next examined the relative effects of alpha power and phase within our pulsed-inhibition model of perception. A key aspect of the pulsed-inhibition model is that the strength of phasic perceptual modulation depends on power [22]. To capture this in our signal detection modelling, alpha power and phase pair-values are mapped onto a single inhibitory value that is used to predict latent sensory modulation on each trial. The mapping to inhibition is shown in the upper panel of Fig. 5a and is increasing with power and pulsed across phase values, consistent with the pulsed-inhibition hypothesis [15, 16]. To examine the dependence of phase on power, we included an additional weighting parameter in the behavioural modelling that describes how much phase contributes to the inhibitory values used to predict latent sensory modulation (fitted freely during the above SDT model inferences). Fig. 5a (lower panel) shows the posterior probabilities for this parameter and reveals that the contribution of phase to behavioural modulation was relatively weak in both the detection (mean weight [95% HDI] = 34% [6.4, 66.2]) and discrimination task (mean weight = 28.3% [2.1, 60.1]). We further visualised the inferred profile of the pulsed-inhibition of response gain in both tasks (see Fig. 5b–c). This shows that the fitted model supports a sustained suppressive effect at high power that is not fully released at less inhibited phases, as pulsed-inhibition theories suggest it should. Together, these analyses suggest that alpha power produces a sustained suppressive effect on behaviour, alongside weaker phasic modulation that does not return to unsuppressed behavioural performance across the phase cycle.

**Figure 5:**
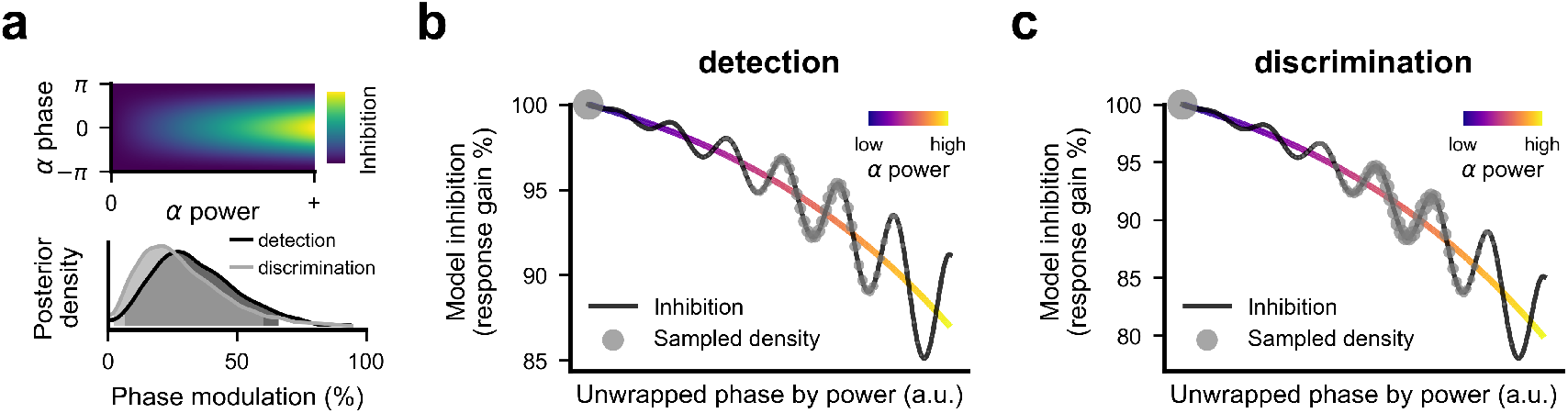
Modelled profile of behavioural inhibition via power and phase. **a** Upper panel: Alpha power and phase values are mapped to an inhibitory value, shown as a two-dimensional space. Lighter colour reflects stronger model inhibition that grows with higher alpha power (x-axis) and is dependent on phase (y-axis). Lower panel: The posterior density of a parameter controlling the percentage extent of phasic modulation. Zero values reflect no contribution of phase corresponding to power modulation only. **b–c** Reconstructed response gain modulation in the detection (**b**) and discrimination (**c**) tasks. The black line shows the model-predicted change in response gain (y-axis) plotted at a continuously unwrapped alpha phase and alpha power (coloured line). Note this relationship is idealised as not all alpha phase-power combinations are shown, but the inhibitory profile is conveyed, nonetheless.

## 3 Discussion

We investigated how human visual sensitivity is shaped by spontaneous alpha-band neural oscillations. Using measures of power and phase at the onset of visual stimulation, we quantified their influence upon psychometric performance in both detection and discrimination tasks. By developing a novel modelling approach, we were able to link power and phase into a combined quantity that reflects an ‘inhibitory drive’ in line with predictions arising from pulsed-inhibition theories [15, 16]. We then inferred the trial-by-trial effects of inhibitory drive upon latent sensory responses within a signal detection theoretic paradigm. Our results show that this combined alpha power–phase predictor is associated with the suppression of sensory evidence, shaping behavioural performance across the full range of presented contrast intensities. Moreover, we uncover the computational basis for oscillatory perceptual effects on sensitivity, finding a response gain mechanism that covaried with alpha oscillations in both detection and discrimination tasks. We further demonstrate that the contribution of phase to the modulation of perceptual performance is far weaker than would be expected from a strong version of the pulsed-inhibition model. Specifically, when alpha power is high, sensory responses are reduced and rhythmically modulated by alpha phase, but there is no phasic return to the unsuppressed levels of behaviour seen when alpha power is low. These results suggest that alpha-frequency neural inhibition manifests in both sustained and weakly phasic suppression of response gain that, together, shape perceptual ability.

A critical feature of our modelling is that it accounts for the variable influence of phase that would be expected at different levels of oscillatory power. That is, the same phase values can be associated with different levels of inhibition depending on power, such that a trough at low power may be associated with perceptual effects that are equal to those of a peak at higher power (see Fig. 5). This dependence of phase effects on power could result in undetectable phase-modulation if not deconfounded. Such a fact has been ignored in most previous investigations of oscillatory phase-effects and may be a reason why some past work (e.g., the contrast binning analyses of [29]; recapitulated here, with results shown in Fig. 3) has apparently supported power-but not phase-modulation of perception. Studies that have acknowledged this issue have typically split their data into binary, high versus low alpha power conditions (e.g., [21, 22, 34]), expecting to observe phase effects only in the high-power condition. Our analysis is robust to these problems, as we model a continuous interactive power–phase relationship and can include all trials during model estimation. This approach can disentangle the possibly opposing effects of phase at different levels of power with no inherent loss of statistical power. More importantly, our combined model of alpha-based inhibition is coherent with theoretical proposals concerning the role of alpha oscillations in neural sensory processing [13, 15, 16]. These theories suggest that the influence of phase depends intrinsically on oscillatory power, due to their common relation via a pulsed inhibitory effect upon neural activity.

It is well established that alpha oscillations inhibit neural spiking [10, 11], but the computational and perceptual consequences of this suppression are currently unclear. Much prior work investigating links between alpha and perception has been unable to speak to the mechanisms involved as it focussed exclusively on hit rates in detection tasks (e.g. [17, 18, 35]). One challenge has been the need to disentangle the influence of alpha on perceptual sensitivity from changes in bias (which may have either perceptual or decisional sources; for review, see [36]). This requires probing behavioural modulation using target-absent trials to allow estimation of the false-alarm rate. Studies employing this approach have typically found that alpha power modulates false alarm rates to the same extent as it modulates successful threshold-level target detections, suggesting a change in the criterion for detection [24, 26, 33]. This approach presents another challenge, however, as performance may be modulated differentially outside the contrast range used to probe threshold-level performance. Studies that have modelled performance over the full contrast range have yielded conflicting results. Chaumon & Busch [29] concluded that alpha power modulation was associated with changes in response gain. In contrast, Pilipenko & Samaha [30] observed a vertical shift in the contrast response function with alpha power that they interpreted as evidence of a change in criterion. However, neither of these studies employed SDT in their modelling of the complete contrast psychometric function and instead infer changes in the psychometric function using a hit-rate axis, which can misrepresent nonlinear effects on sensitivity. By estimating the source of alpha modulation across varying levels of contrast in a latent signal-detection model, we were able to show that perceptual sensitivity is affected in a manner consistent with changes in response gain across both discrimination and detection tasks. This suggests that the inhibition associated with alpha oscillations divisively scales neural responses depending on the strength of sensory input and thus alters perceptual sensitivity.

In addition to response gain modulation in both tasks, we observed a robust alpha-related change in criterion that was unique to the detection task. This result is consistent with past findings [24, 26, 33] and suggests that alpha-frequency inhibition modulates baseline firing rates in visual cortex [36]. This modulation of baseline firing alters the probability of reporting a stimulus as present on the basis of noise excitability in the visual system (i.e. false alarms). It will, however, have little influence on discrimination performance, if perceptual noise is equally likely to affect both response options, and so result in no overall change in bias [26]. In line with this, we did not observe a criterion effect in our discrimination task. Together, the effects of response gain and criterion suggest a mechanism by which alpha-based inhibition scales visual neural activity without altering underlying sensitivity to contrast (i.e. via a threshold or contrast gain change). This would be consistent with inhibition of visual cortex that scales all components of cortical firing activity, including both baseline neural activity and the afferent stimulus-driven responses from earlier neurons.

Our findings suggest that over the cycle of the alpha oscillation, alpha phase is associated with similar mechanistic fluctuations as alpha amplitude. Critically, this was true across both tasks. These results provide a mechanistic explanation for previous findings showing fluctuations of detection across the alpha cycle [17, 18, 21, 37, 38], and align with recent results that indirectly suggest alpha phase and amplitude are associated with the perceptual consequences of lateral inhibition in early visual regions [39]. Importantly, we also observed direct evidence for suppression of visual responses as alpha power and phase both reduced the magnitude of stimulus-evoked neural response (see also, [21, 33, 37, 40]). These results provide converging evidence for the suppression of early visual responses by alpha-frequency neural oscillations (c.f., [41]).

Our pulsed-inhibition model is a direct implementation of the theorised relationship between alpha-frequency oscillations and the underlying neural activity that is relevant for perceptual processing [13–16]. Signal detection modelling also allowed us to compare the relative contributions of alpha power and phase toward the modulation of perception. Theory suggests these should be equivalent; the effect of power should simply be the effects of phase, at that power level, integrated over the course of an oscillatory cycle. However, we observed that the relative effect of phase in this model was only ∼30% of what theories suggest it should be, across both tasks. Such a finding may provide a plausible explanation for why phase effects have been difficult to observe in some scenarios (e.g. [34]; see also [42]). The source of this discrepancy is difficult to pinpoint. Electrophysiological features that were not included in our modelling, such as the non-sinusoidality of posterior alpha [43] or saturation of modulation at much higher power levels, could weaken an observed phase-power correspondence. However, our results suggest the weakened influence of phase was present even at moderate levels of power, so it would be surprising for either of these possibilities to reduce the observable phase effect by ∼70%. Instead, our results suggest the existence of a tonic, non-pulsatile suppressive effect that scales with alpha amplitude. One candidate mechanism are GABAergic inhibitory networks that are known to be associated with alpha oscillations (e.g. [44]; [45]) and commonly have time-constants longer than the ∼100 ms cycle of an alpha oscillation. The activation of GABA-B receptors in visual cortex (and elsewhere) can produce inhibitory effects spanning hundreds of milliseconds [46], suggesting a possible mechanism by which alpha oscillations could produce tonic inhibition of visual processing.

In summary, here we used a model-based analysis of visual behaviour to reveal the mechanistic bases by which measured neural oscillations in the alpha band shape perceptual sensitivity. Our results suggest that sensory evidence is divisively suppressed, in line with the idea that alpha oscillations reflect an inhibitory influence upon neural processing. We show that the strength of phasic modulation is not fully consistent with a strong version of the pulsed-inhibition model. Rather, our results suggest the effects of alpha phase may be weaker than the largely tonic effects of alpha power on perceptual behaviour. These findings underscore the necessity of modelling brain activity and perception jointly to explain the nonlinear interactions that link alpha-band dynamics to perceptual behaviour.

## 4 Methods

### 4.1 Participants

Nine healthy adults with normal or corrected-to-normal vision participated in the study (mean age, 26.9 years; range, 22–40 years; five females), including two non-naive authors HB and AH. Experiment 1 (detection task) involved eight participants (4 males and 4 females). Experiment 2 (discrimination task) involved the same participants, but one female dropped out and was replaced by another female participant. No participants were excluded from any analyses. The experiment was approved by The University of Queensland Human Research Ethics Committee. All participants gave written informed consent and all but the authors were compensated at a rate of $20/h AUD.

### 4.2 Apparatus

The stimuli were presented on a gamma-corrected 24-inch LCD monitor (VIEWPixx 3D, VPixx Technologies) with 1920 x 1080 display resolution and 120 Hz refresh rate within a dark, electromagnetically shielded room. Viewing distance was maintained at 58 cm using a chin rest. Stimuli were generated using MATLAB (MathWorks, R2020a) with the Psychophysics Toolbox v.3.0.17 [47, 48]. A Biosemi ActiveTwo system (Biosemi, Amsterdam, Netherlands) was used to record 64 Ag-AgCl electrodes digitised at 1024 Hz and arranged in the standard 10-10 layout [49]. Per Biosemi design, the Common Mode Sense and Driven Right Leg electrodes served as reference and ground. Eye muscle activity was monitored using two EOG electrodes placed above and below the right eye, and two placed at the outer canthi of each eye.

### 4.3 Stimuli, task, and experimental procedure

The stimuli were sinewave gratings (4 cpd; ±45^*°*^ random orientation; randomised phase) presented within a Gaussian window (0.5 dva sigma) on a mid-grey background. A centrally positioned black cross (size: 0.2 *×* 0.2 dva; line width: 2 pixels) was presented throughout the trial to reduce eye movements. The target stimulus was presented at a single lower left location on all trials (7.07 dva eccentricity, 5 dva below and 5 dva to the left of fixation) to maximise trial numbers for analyses without needing to counterbalance analyses across hemispheres. In the detection task (experiment 1) the stimulus was presented within two spatial markers (line lengths: 0.2 dva; line width: 2 pixels) positioned along the 45^*°*^ diagonal and offset from the centre of the stimulus by 2 dva (4 *×* sigma). The observers’ task was to report whether the target stimulus was present or absent on each trial. In the spatial discrimination task (experiment 2), the stimulus was presented 1 dva along the opposite diagonal perpendicular to the spatial markers, and the observers task was to report whether the stimulus appeared to the left or right of the spatial markers (Fig. 2a).

Each trial started with a variable fixation period of 1500–2000 ms. The stimulus was then presented for 8.33 ms and observers had 1500 ms to make a response. In the detection experiment, the colour of the fixation cross was changed from black to blue at the stimulus onset, to remove temporal uncertainty. In the discrimination experiment, the fixation colour change occurred 1500 ms after the stimulus, to inform the participant that the stimulus was no longer displayed and a two-alternative forced choice response was required. Participants completed an hour-long introductory behavioural testing session to gain familiarity with each task prior to EEG recordings. Each participant then completed four EEG sessions of approximately 400 trials, resulting in 29,090 trials for analysis in experiment 1 and 22,020 trials in experiment 2. Breaks were given every few minutes.

Psychometric functions were fit online using a Bayesian approach to continually update a parameter posterior using a discrete grid approximation. We used a cumulative Gaussian psychometric function with parameters for the slope and location in log contrast, as well as independent upper and lower asymptotes. On the first 150 trials in each session, and a random 10% of trials thereafter, the stimulus contrast was adaptively determined using a look-ahead algorithm to find the contrast value that minimised the expected entropy of the parameter posterior [50]. This allowed us to present contrast values that were most informative for quick estimation of the psychometric function parameters, and to ensure that this estimate continued to be correct throughout the session, as needed. On the remaining (majority) of trials, contrast was sampled from a normal distribution in log contrast centred on the current best estimate of the participants’ threshold and with standard deviation of the inverse slope estimate. This ensured that the majority of stimulus values were sampled around the most informative parts of the psychometric function, where performance changes rapidly with contrast [51]. In the detection experiment, an additional random 10% of the non-adaptive trials were chosen for zero-contrast stimuli to enforce extra sampling of the false alarm rate for signal detection analyses.

### 4.4 Electroencephalography

#### 4.4.1 Recording and preprocessing

Offline processing of EEG data was performed using EEGLAB v2024.2 [52]. Signals were downsampled to 512 Hz, high-pass filtered at 0.5 Hz using *pop_eegfiltnew*.*m*, and re-referenced to the average of electrodes. Bad channels were automatically identified using the adaptive channel scoring algorithms from the FASTER pipeline [53] and were interpolated using spherical splines. Independent component analysis (ICA) was then performed with the extended Infomax algorithm [54] using *pop_runica*.*m* on a copy of the data that was 1 Hz high-pass filtered, demeaned across epochs [55], and automatically cleaned of noisy epochs using the *pop_autoreg*.*m* function with default parameters. Fitted ICA weights were then used to project the original data and epochs into the component space.

We used a semi-automated procedure to select focal components for further analyses. First, all components were ranked according to the unsigned correlation between the component activation maps and a posterior region of interest contralateral to the stimulus (electrodes Oz, O2, POz, PO4, PO8, P2, & P4). We then selected a single component from the top five ranking components. This was done by manually inspecting the trial-averaged prestimulus spectral density and the poststimulus evoked waveform for each component (see Supplementary Figures 2 and 3 for the component topographies, evoked responses, and frequency spectra of chosen components). The highest ranking component (commonly the first or second) that showed both an alpha spectral peak and a typical visually-evoked response was selected for analysis. To enforce sign consistency across participants and sessions, the sign of the component was flipped so that the maximally activated electrode was positive (this also ensured a common oscillatory phase interpretation across participants).

#### 4.4.2 Oscillations

Prestimulus oscillatory power was estimated using a Fourier transform on the hamming windowed data within a −0.5–0 s prestimulus epoch (2048 sample padding). The *specparam* toolbox [32] was then used to separate aperiodic and non-aperiodic components within the power spectra. Frequencies between 1–40 Hz were used with the algorithm’s default parameters, maximum peaks set to 4, and peak width limits set between 0.5–6 Hz (trial-averaged *R*^2^ = 0.81). From this, we extracted the peak frequency of detected oscillations within the 8–14 Hz frequency range at the single trial level. To then characterise alpha power and phase, we used the endpoint-corrected Hilbert transform method [56] to estimate the instantaneous power and phase at stimulus onset. We applied this only on trials in which *specparam* detected an oscillation. This was done by band-pass filtering prestimulus epochs (−1.5–0 s) between +/-1 Hz of the peak frequency, and then using the absolute value and angle of the resulting complex analytic signal as power and phase estimates, respectively.

#### 4.4.3 Analyses of evoked responses

To characterise the magnitude of evoked responses, we detected the timepoint at which the (negative) peak minima was found in the poststimulus data. This was done separately per session using the averaged waveform of all trials in that session, and searching within a 160–240 ms window. To yield a single-trial measure of evoked amplitude, trial-level data were averaged within a 30 ms time window centered on the per-session peak timepoint. The time window used for automatic selection was determined by inspecting the waveform grand-averaged across all data.

A mixed effects General Linear Model (GLM) was used to predict the evoked response magnitude using alpha power. Random effects were fit for the intercept and slope terms. The effect of alpha phase was analysed using separate linear regressions, per session, of the sin and cos of phase. Session-specific analyses allowed the optimal phase of the regressed sinusoidal modulation effect to vary between participants and sessions, accounting for potential phase differences that stem from individual cortical differences and electrode placements in each session. We assessed group level significance by combining *P* values using Fisher’s method. An effect size, *d*_*z*_, was calculated for group-level phasic modulation by reconstructing the amplitude of phasic modulation from regression weights, and computing a *z*-score across sessions.

### 4.5 Behavioural analyses

#### 4.5.1 Data transformation and normalisation

Contrast, power, and phase data were normalised across sessions, allowing data to be collapsed together, while mitigating major subject-specific differences. Contrast was computed as a proportion of the threshold contrast that was adaptively located during each session. Alpha power was log transformed and mean-scaled, whereby non-zero power values were divided by the average power, per session, to express power in relative units (mean = 1). Given that we found significant group-level phasic modulation of the evoked response, we took the specific phase of these effects as the relevant phase for potential behavioural modulation. Thus, alpha phase was normalised by circularly subtracting the phase value that predicted the strongest evoked response magnitude, specific to each session. For analyses that model a linear phase effect (specifically, binning and logistic regression) phase was transformed by taking the absolute value, which yields an unsigned distance from the optimal evoked phase. Trials containing no detected alpha oscillation were assigned a normalised power value of zero and were also excluded from analyses that modelled phase exclusively (as phase is not meaningful in zero-power oscillations).

#### 4.5.2 Binning analyses

We used eight equally sized contrast bins, combining data across participants. Within each contrast bin the relevant alpha data (either power or phase) were further divided into four equally sized bins. Behavioural responses were then averaged within each contrast-power/phase bin. A first degree polynomial was then fit across the alpha power/phase bins to yield the slope of accuracy modulation within each contrast bin. We then took the average slope over contrast bins as an aggregate measure of behavioural modulation across contrast bins. This procedure was repeated 5,000 times using bootstrapped datasets. We calculated bootstrapped *P* values using the one-sided probability that samples lay above zero. That is, under the null hypothesis that alpha power/phase suppress behavioural performance, we would expect the bootstrapped aggregate slope samples to lie at or above zero.

In the discrimination task, behavioural choices were modelled as a function of signed contrast, using the difference in contrast between the right and left stimulus positions (Fig. 3b). This results in better accuracy when the probability of reporting the righthand position is higher for positive signed contrast and lower for negative signed contrast. We accounted for this by inverting the slopes of negatively signed contrast bins prior to aggregating across contrast bins. We included zero-contrast trials in an additional bin that was excluded when calculating the aggregate slope, as performance here is at chance level by definition (i.e. guessing).

#### 4.5.3 Logistic regression

Generalised Linear Mixed Effects models were used to assess the influence of alpha power and phase on behavioural accuracy, using MATLAB’s *fitglme* with a logistic link function. Model fit was improved by including predictors for additional transformations of contrast, by raising contrast to exponents of 0.5 and 2. Including these terms models the nonlinear relationship between contrast and predicted behavioural accuracy, which is achieved by approximating curved changes on the linear predictor scale instead of utilising only the sigmoidal shape of the logistic link function [57]. We included random intercepts in the model of the contrast terms and use this as a null model that fitted performance on the basis of stimulus contrast alone. Additional models were fitted using alpha power and phase, including intercepts for these terms as well as slopes that interacted with the contrast terms and thus describe how power and phase shape a contrast-related change in accuracy. Finally, we fit a power by phase interaction model. The significance of predictors was assessed using nested likelihood ratio tests to assess the contribution of increasing model complexity. This allowed us to infer the significance of power, phase, and power by phase interaction terms, by looking across the additional terms that interact with contrast. We also evaluated model performance by transforming information criteria (AIC, BIC) into weighted model probabilities (see Supplementary Figure 1; [58]).

### 4.6 Signal detection modelling

We used a signal detection theory approach to model behavioural responses in both tasks. The stimulus was assumed to be represented in a one-dimensional sensory space, and its representational intensity varied on a trial-by-trial basis. This variation was modelled as a unit variance Gaussian distribution where the mean stimulus representation intensity is a function of the external stimulus contrast:

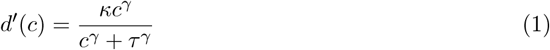

This function describes a saturating non-linear mapping of the stimulus contrast, *c*, onto an internal response strength (i.e. a *transducer function*) that has been studied in neurophysiology [27, 59, 60]. The parameter *κ* controls the maximum strength of internal representation that reaches saturation, *τ* locates the contrast at which the function becomes compressive and begins to saturate, and *γ* controls the rate at which stimulus contrast increases the internal response. In the detection task, it was assumed that the observer responds with a yes decision on trials where this internal representation exceeds a fixed criterion, *λ*. A decision rule for the localisation task is explained further below. The probability of a binary yes-no decision is thus given by integrating the probability that the internal representation exceeds *λ* for a given stimulus contrast:

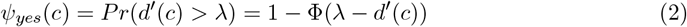

where Φ(·) is the cumulative Gaussian function, which performs an identical role to the probit link function within the General Linear Model [61]. The observed detection responses are modelled as a Bernoulli random variable where the probability of success is a given by the transducer function of contrast:

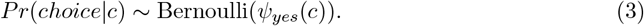

In the discrimination task, the observer chooses the spatial location of a single target stimulus (left or right of the spatial markers). We modelled this task as a 2-alternative forced choice (2AFC), in which the observer monitors both spatial positions and reports the location with the higher internal response on each trial. The probability of reporting the right-hand location is:

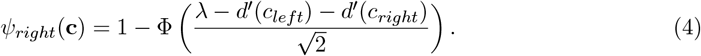

In this context, the criterion *λ* is a bias parameter that shifts the decision in favour of a certain spatial interval but does not reflect a subjective threshold for the stimulus representations that are compared objectively. Note that we modelled the choice of a single response option, which ranged from 0–100% correct rather than 50–100% correct used for accuracy [62]. We also modelled an additional lapse rate parameter in the discrimination task, which improved model fit and accounts for the larger number of guessing responses caused by additional temporal uncertainty in the task design. With lapse proportion, *ϵ*, the corrected response probability is 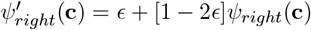.

#### 4.6.1 Pulsed-inhibition modelling

To combine alpha power, *P*, and phase, *ϕ*, values into a single scalar inhibitory value, *I*, we used the function on each trial, *i*:

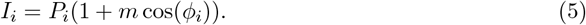

This mapped single-trial power and phase into a value that was used as a covariate within the signal detection modelling. The weighting parameter *m* ∈ [0, 1] controlled the extent of phasic modulation, with zero values providing no phasic modulation (i.e. a power-only modulation effect) and a value of one providing full phasic modulation (consistent with the functional form visualised in Fig. 1a). We tested four candidate mechanisms by which inhibition could modulate sensory processing. For response gain modulation, the output of the transducer function was scaled, using a log link function for inhibition modulation:

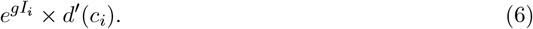

Here, the free parameter *g* controls the predicted modulation given inhibition and contrast values. Note that multiplying the output of the transducer function is mathematically equal to adjusting the *κ* parameter in 1. For contrast gain modulation, the transducer function threshold parameter, *τ*, was adjusted using a logit link for inhibition:

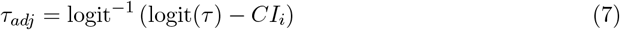

where *C* is free parameter controlling contrast gain modulation. For noise modulation, the probability of task responses was modulated by divisively scaling the input to the cumulative Gaussian, *G*, in both *ψ*_*yes*_ and *ψ*_*right*_ defining

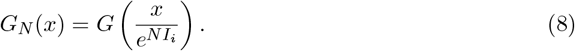

Here, *N* is the free parameter controlling noise modulation via a log link for inhibition. Criterion modulation was achieved with the free parameter, *b*, by adjusting *λ*:

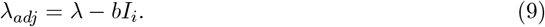

We fit the Bernoulli response models described above to the single-trial data within a Bayesian framework using Hamiltonian Monte Carlo in PyMC v4 [63]. Parameters were modelled using a mixed-effects approach to hierarchically model the variation across the 32 experimental sessions for each task (individual psychometric function fits are shown in Supplementary Figure 4). Here, participant-level contrast thresholds, *τ*, were fitted, so the untransformed contrast values were modelled. As *d*^*′*^ is difficult to estimate at high values, we followed Lesmes *et al*. [62] and fixed the parameter *κ* at 5. Before fitting the observed data, we conducted a prior predictive analysis to ensure that the choice of priors was able to generate model predictions that were psychophysically plausible and consistent *a priori* with realistic data (i.e. monotonically increasing psychometric functions, without extreme/flat slopes, and without contrast thresholds near full/zero contrast). For the *m* parameter controlling phasic modulation, we used a flat Beta(1,1) prior. For the remaining modulation priors (*b, g, N, C*), we used weakly regularising Normal(0,2) priors.

#### 4.6.2 Statistical inference

Models were fit using the No U-Turn Sampling HMC method in PyMC. 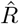 values were less than 1.01 for all parameters, suggesting that sampling chains converged to the target posterior. The reported *β*_*z*_ values in text and figures refer to the standardised modulation parameters. These were obtained by dividing the posterior samples of the group-level modulation parameters (*b, g, N, C*) by their fitted standard deviations. Models were fit independently to facilitate interpretation, as parameters within the probit link are highly dependent on the specific values of all other parameters. For the detection modelling we verified that support for the response gain and criterion results remained when fit together, and we report this combined model in text and figures.

We conducted hypothesis tests using the inferred posterior probabilities of the standardised effects, and we assessed whether the 95% Highest Density Interval (HDI) of the posterior contained the null value (zero). We also performed Bayes Factor analyses to assess the strength of evidence for modulation. The fitted hierarchical models gave us insight into the variability of modulation effects (which we modelled as normally distributed across sessions), and this also provides information on the parameter value at the group level. We computed Bayes Factors using the posterior mean estimates of the standardised modulation effects. This was calculated on the basis that the standardised effect is expected to be zero under the null hypothesis (i.e. there is no effect relative to residual variability in the parameter) and follows a Cauchy distribution (with standard scale parameter of 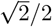, [64]) under the alternative hypothesis.

## 5 Data availability

The behavioural and neural data generated in this study are available on the Open Science Framework database: https://osf.io/3rm8z/.

## 6 Code availability

Code has been deposited on GitHub: https://github.com/henrybeale/alpha-power-phase-visual-sensitivity.

## 7 Acknowledgements

AH was supported by the Australian Research Council (DE2200101019). JM was supported by a National Health and Medical Research Council (Australia) Investigator Grant (GNT2010141).

## 8 Author contributions

HB contributed to the conceptualisation, methodology, software, formal analysis, investigation, data collection, data curation, visualisation, and writing – original draft and revisions. JB contributed to conceptualisation and writing – revisions. AH contributed to conceptualisation, methodology, investigation, project administration, and writing – original draft and revisions.

