## Supplementary Figures for "A joint alpha power–phase dynamic shapes visual sensitivity"

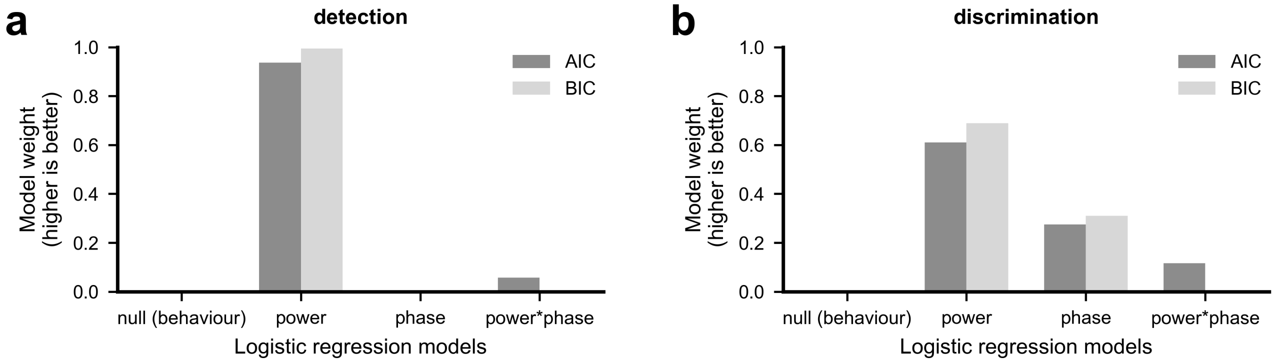


**Figure S1:** Model weights for logistic regression models. Weights were created using AIC (dark bars) and BIC (light bars).

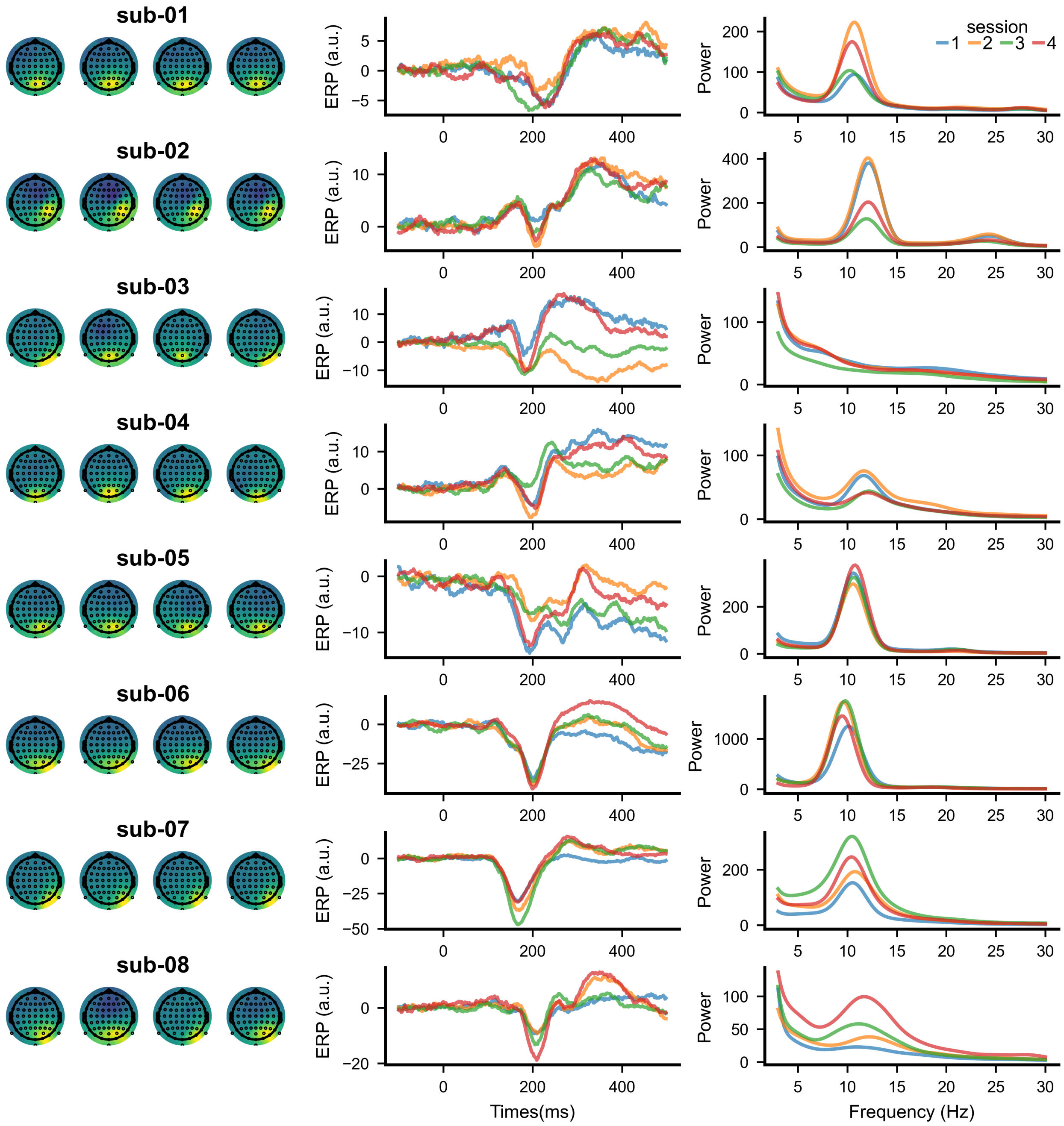


**Figure S2:** EEG topographies (left column), evoked responses (middle column), and prestimulus frequency spectra (right column) for the detection task. Topographies are z-scored activation maps of the selected independent component for each session. The frequency spectra were decomposed using the -500–0 ms stimulus-locked component activation (using a hamming window).

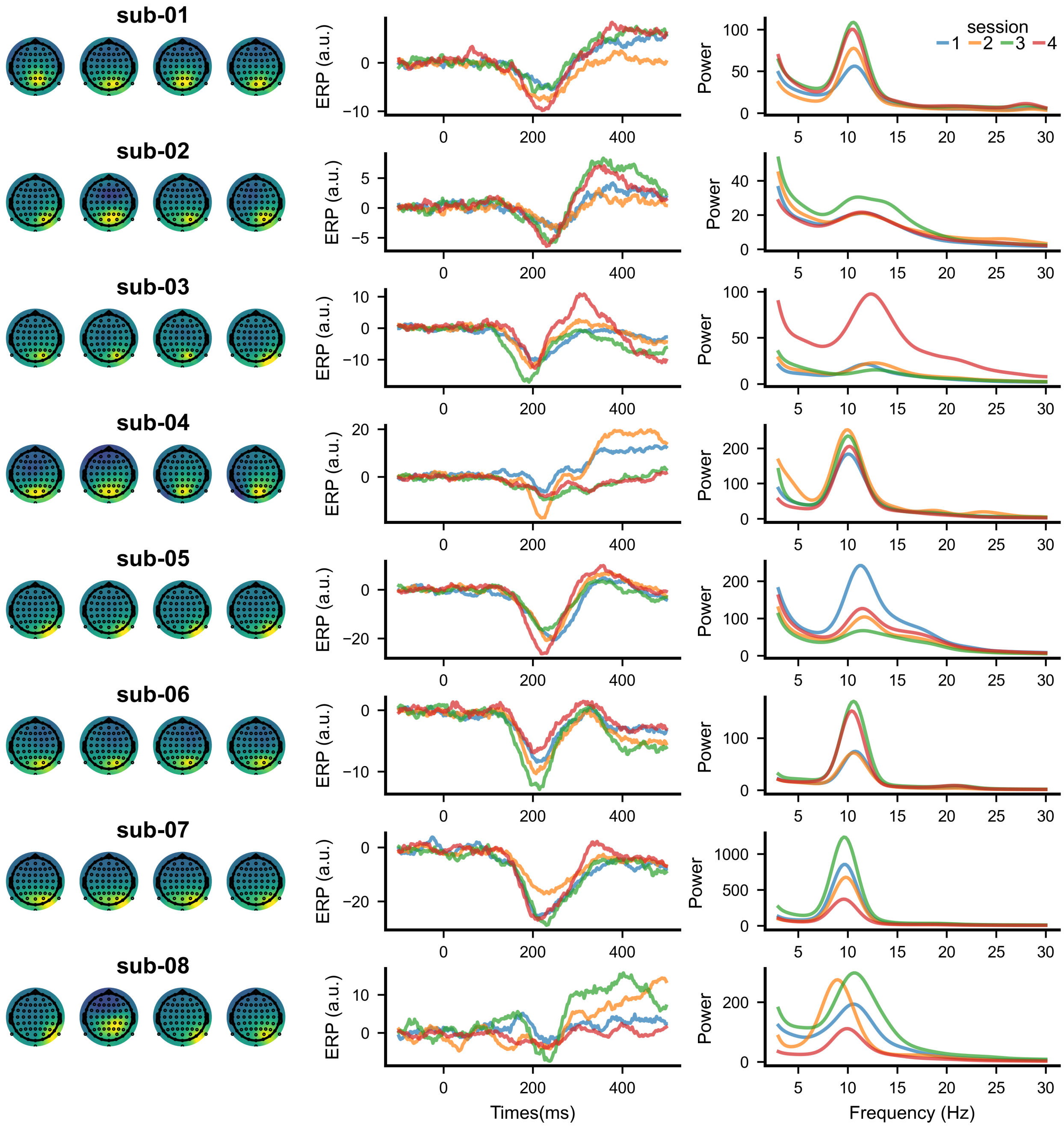


**Figure S3:** EEG topographies (left column), evoked responses (middle column), and prestimulus frequency spectra (right column) for the discrimination task.

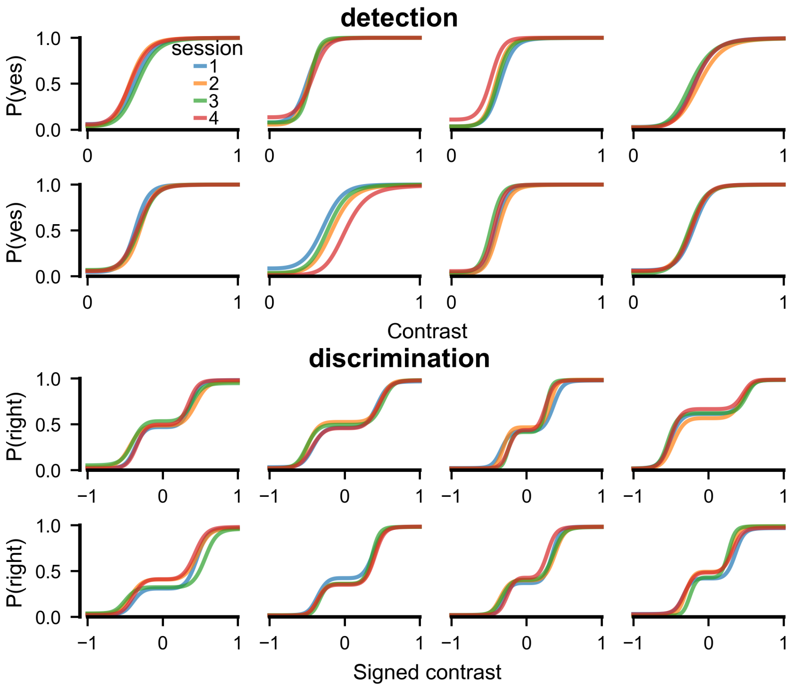


**Figure S4:** Psychometric function plotted for eight observers (shown one per panel) over their four sessions (coloured lines) of the detection (top) and discrimination task (bottom). The functions were fit hierarchically to account for group level variation between sessions.
